# Plant-produced RBD and cocktail-based vaccine candidates are highly effective against SARS-CoV-2, independently of its emerging variants

**DOI:** 10.1101/2022.04.07.487347

**Authors:** Tarlan Mamedov, Damla Yuksel, Irem Gürbüzaslan, Merve Ilgın, Burcu Gulec, Gulshan Mammadova, Aykut Ozdarendeli, Shaikh Terkis Islam Pavel, Hazel Yetiskin, Busra Kaplan, Muhammet Ali Uygut, Gulnara Hasanova

## Abstract

SARS-CoV-2 is a novel and highly pathogenic coronavirus, which has caused an outbreak in Wuhan City, China, in 2019 and then spread rapidly throughout the world. Although several COVID-19 vaccines are currently available for mass immunization, they are less effective against emerging SARS-CoV-2 variants, especially the Omicron (B.1.1.529). Recently, we successfully produced receptor-binding domain (RBD) variants of spike (S) protein of SARC-CoV-2 and an antigen cocktail in *Nicotiana benthamiana*, which are highly produced in plants and elicited high-titer antibodies with potent neutralizing activity against SARS-CoV-2. In this study, we demonstrate that these protein-based vaccine candidates are highly effective against Delta and Omicron variants. These data support that plant produced RBD and cocktail-based antigens are most promising vaccine candidates and may protect against Delta and Omicron-mediated COVID-19. Based on the neutralization ability, plant produced RBD and cocktail-based vaccine candidates are highly effective against SARS-CoV-2, independently of its emerging variants.

## Introduction

The SARS-CoV-2 virus that causes the COVID-19 infection, a highly infectious RNA virus, has undergone numerous mutations with the formation of genetically diverse linkages since first appearing in the city of Wuhan in 2019. Mutations affecting the interaction between the S protein and the ACE-2 receptor, potentially affect the viral transmissibility. Among variants of concern (VOCs), the Alpha (B.1.1.7) and Delta (B.1.617.2) variants are associated with a high viral transmission rate and virulence compared to the parental Wuhan strain. Nine mutations were found in the S protein of the Delta variant, including in the amino terminal domain (NTD) (five mutations) and RBD (two mutations, L452R, T478K). One mutation was found to be close to the furin cleavage site (P681R), and one in the S2 region (D950N) (*1*). Notably, mutations of G476, F486, T500, and N50, observed within the RBD are close to the ACE-2, a receptor binding site.

On 26 November 2021, a new variant named Omicron (B.1.1.529) was designated the fifth VOC (*2*), which carries more that 60 mutations compared to the original Wuhan strain. In the new Omicron variant, 36 mutations were found in the S protein sequence, including 30 amino acid substitutions, 3 deletions and 1 insertion. It should be noted that 15 of the 30 amino acid substitutions are in the RBD (*3*).

Vaccination is currently the most effective way to prevent from pathogenic diseases, including COVID-19. However, currently available vaccines are less effective or not effective against emerging SARS-CoV-2 variants, such as the Delta strain (B.1.617) and the Omicron (B.1.1.529). Therefore, developing COVID-19 vaccines that are effective against emerging new variants of SARS-CoV-2, such as Omicron, will be a very challenging task. Recently, we reported the successfully production of RBD variants of S protein of SARS-CoV-2 (*4*) and of an antigen cocktail (*5*) in *Nicotiana benthamiana*, which are highly produced in plants and elicited high-titer antibodies with potent neutralizing activity against SARS-CoV-2. In this study, we demonstrate that plant produced RBD or cocktail antigen-elicited antibodies are capable of neutralizing the Delta or Omicron variants.

## Methods

### Cloning, expression and purification of gRBD, dRBD and N+RBD proteins

Cloning, expression and purification of gRBD, dRBD and N+RBD proteins from *N. benthamiana* plant were performed as decribed recently (4,5). Purification of plant-produced gRBD, dRBD and N+RBD proteins was performed from 20 g of frozen leaves, infiltrated with the pEAQ-RBD-Flag-KDEL (with or without pGreenII-Endo H), or pEAQ-RBD-Flag-KDEL+ pEAQ-N-Flag-KDEL constructs, using anti-FLAG affinity chromatography as described recently (4,5).

### Immunogenicity Studies

Immunogenicity studies of gRBD, dRBD and N+RBD in mice were perforemed in groups of 6–7 week old Balb/c male animals (six mice/group) as described recently (4). Mice were immunized intramuscularly (IM) on days 0 and 21 with 5 μg of gRBD, dRBD and N+RBD adsorbed to 0.3% Alhydrogel. Blood samples were taken from immunized mice on days 42 used for microneutralization assay (MNT). Mice studies were conducted at Akdeniz University Experimental Animal Care in compliance with the ARRIVE guidelines, and with the permission of the Animal Experiments Local Ethics Committee for Animal Experiments at Akdeniz (under protocol number of 1155/2020.07.0) with the supervision of a veterinarian.

### MNT assay

We used to live the SARS-CoV-2 Wuhan (GB-MT327745; GISAID-EPI_ISL_424366), Delta (GB-OM945721;GISAID-EPI_ISL_10844545), and Omicron variants (GB-OM945722; GISAID-EPI_ISL_10844681). SARS-CoV-2 microneutralization (MN) tests were performed as previously described with minor modifications (4). Before the day of the experiment, Vero E6 cells (ATCC, CRL-1586) (2×104 cells/100ul/well) were passaged to 96 well plate. The sera collected from vaccinated mice were heat-inactivated for 30 min at 56°C and was subjected to two-fold serial dilutions (from 1:4 to 1:1024) with serum free Dulbecco’s modified Eagle’s medium (DMEM). Two-fold serial dilutions of the mice sera were mixed with an equal volume of DMEM containing 100 tissue culture infectious dose 50 (100 TCID50) of the SARS-CoV-2 variants and incubated at for 90 min at 37°C. Following adsorption, inoculums were removed and cells were incubated for 72 h with DMEM containing 2% FBS and checked for the cytopathic effect (CPE). The microplates were designed to test two wells for each serum dilution, six virus-only control wells, and three blank wells containing growth medium alone. The CPE evaluation was performed according to the reduction of infection by 50% or more at the serum dilutions. The 50% microneutralization titre (MNT50) was analysed by the Spearman-Karber method in which calculated as the reciprocal of the highest serum dilution at which the infectivity was neutralized in 50% of the cell in wells.

## Results and discussion

To evaluate whether RBD or cocktail antigen-elicited antibodies are capable of neutralizing the Delta or Omicron variants, the latter being dominant at the moment, we tested Wuhan (GB-MT327745; GISAID-EPI_ISL_424366), Delta (GB-OM945721; GISAID-EPI_ISL_10844545), or Omicron variants (GB-OM945722; GISAID-EPI_ISL_10844681) with sera of mice immunized with two doses (5 μg per dose) of the plant produced RBD variants or cocktail antigen-based COVID-19 vaccines. After two doses, Delta-neutralizing titers were not significantly reduced compared with Wuhan-neutralizing titers. Under the same conditions, Omicron-neutralizing titers were reduced by not more than 4-fold compared with neutralizing titers of Wuhan (Figure 1). The plant produced, deglycosylated RBD vaccine candidate was more effective compared to its deglycosylated counterpart, suggesting the negative effect of N-glycosylation on protein functionality. It should be noted that, after two doses (Administration of two 30-μg doses of BNT162b2 to participants, with a total of 60μg of synthetic mRNA), the messenger RNA (mRNA)–based COVID-19 vaccine (BNT162b2) was ineffective against Omicron and Omicron-neutralizing titers were reduced by more than 22-fold compared with Wuhan-neutralizing titers (*6*). Cele et al., (2022) demonstrated that in two-dose vaccinated individuals with the BNT162b2, the neutralization protection dropped over 40-fold against the Omicron versus the ancestral D614G virus. These results were expected given that the BNT162b2 vaccine is based on the full-length S protein sequence while the S protein of the Omicron variant is highly mutated compared to other variants (*7*).

**Fig. 1.**
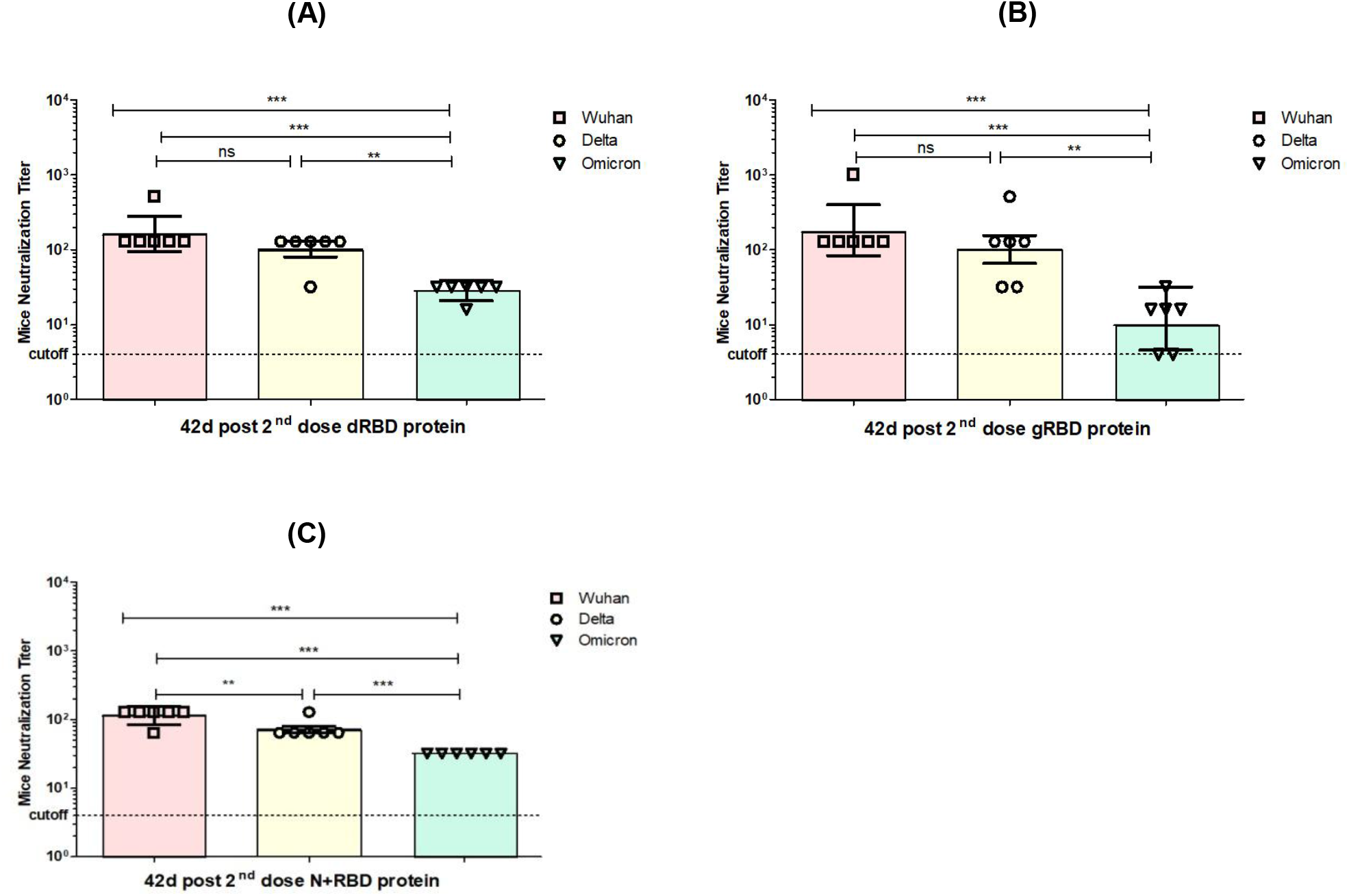
50% microneutralization titers (MNT_50_) of sera from BALB/c mice immunized with plant-produced gRBD, dRBD and N+gRBD(cocktail) proteins against live SARS-CoV-2 the Wuhan, Delta, and Omicron variants. Microneutralization assay of 42nd day mouse sera immunized with plant-produced gRBD, dRBD and N+RBD (~2.7 μg + 2.3 μg, respectively) against live SARS-CoV-2 Wuhan, Delta and Omicron variants as indicated. A: microneutralization of sera from mice immunized with 5 μg of gRBD on day 42; B: microneutralization of sera from mice immunized with 5 μg of dRBD on day 42; C: microneutralization of sera from mice immunized with 5 μg of N+gRBD (~2.7 μg + 2.3 μg, respectively) on day 42. The experiment was performed using 4 to 1024 dilutions of mouse sera collected on day 42. One-way ANOVA Tukey’s multiple comparison tests were used to calculate statistical significance (n = 6 mice/group); *** p < 0.001.

Plant transient expression systems have proven to be promising expression platforms for the expression of variety of important recombinant proteins such as vaccines (*8–12*), therapeutic proteins (*9*), and antibodies, human and industrial enzymes. Importantly, plant expression systems have proven to be capable of high-level production of functional active SARS-CoV-2 proteins (*4,5,13*) and ACE2 (*14,15*), receptor of SARS-CoV-2 and SARS-CoV. We recently demonstrated the successful expression of glycosylated and deglycosylated forms of RBD and antigen cocktails, comprising RBD and nucleocapsid (N) proteins, as promising vaccine candidates against COVID-19. We demonstrated that RBD variants of S protein of SARS-CoV-2 and antigen cocktail in *Nicotiana benthamiana* are highly produced in plants and elicited high-titer antibodies with potent neutralizing activity against SARS-CoV-2. Our hypothesis was that if any SARS-CoV-2 variant infects a human, this means that a mutation in the RBD region did not affect its binding to ACE2, the SARS-CoV-2 S protein receptor. Therefore, the correct selection of RBDs is critical for the successful production of functional RBDs with the ability to induce high neutralizing antibodies against SARS-CoV-2 and its variants. Since SARS-CoV-2 is an mRNA-based virus and mutations were expected, our strategy to eliminate possible emerging mutations was to select not the full sequence of the spike protein (most COVID-19 vaccine developers including Pfizer-Biontech and plant based, Medicago’s VLP, which was approved for use by Health Canada, were targeted on the full-length S sequence), but rather a specific region of the S protein covering the RBD. In addition, to address mutations, the N-protein + RBD multi-antigen vaccine was developed and produced for the first time as a potential COVID-19 vaccine candidate (*5*).

S protein of SARS-CoV-2 is a cysteine-rich protein and nine cysteine residues are found in the RBD and eight of them are involved in the formation of four disulfide bridges (*16*). Notably, for some viruses, such as HIV, the redox state of the fusion protein was shown to be important for the viral fusion to the target cells (*17,18*). Thus, proper formation of disulfide bridges is critical for proper folding of the RBD. We produced RBD variant consisting of amino acids R319-S591, where there is an even number of cysteine residues, which form correct disulfide bridges in the molecule that may stabilize the protein conformation leading to a functional protein. In fact, as we recently reported, the expression levels of both glycosylated (gRBD) and deglycosylated (dRBD) RBD (*4,13*) were greater than 45 mg/kg of fresh weight. The purification yields were ~22 mg of pure protein/kg of plant biomass for gRBD and ~20 mg for dRBD, which would be sufficient for the commercialization of these vaccine candidates. Moreover, in mice, the plant-produced RBD antigens elicited high titers of antibodies with a potent virus-neutralizing activity (*4*). We concluded that the correct selection of the amino acid region of the RBD is crucial for high-yield production of a functionally active and soluble protein (*4*). On this point, in the study of Rattanapisit et al. (2020), RBD of Spike protein of SARS-CoV-2 was produced in *N. benthamiana* plant and low yields (2-4 ug/g of fresh weight) were reported (*19*). In this study the amino acid sequence (F318–C617) of RBD was not properly selected and as a result, cytosine at position 617 remained unpaired (which should form a disulfide bridge with C649 in the full-length S protein), which may destabilize the protein conformation leading to a loss of functional activity. In fact, there was no report of neutralization of the SARS-CoV-2 in this study. In another study, Siriwattananon et al. (2021), F318-C617 amino acids of RBD with Fc region of human immunoglobulin G1 (IgG1) was selected for production in *N. benthamiana* plant (*15*). As in the study of Rattanapisit et al. (2020), the amino acid sequence of RBD (F318–C617) was not properly selected and as a result, cytosine at position 617 remained unpaired (*19*). The expression level of plant-produced SARS-CoV-2 RBD-Fc was low, 25 μg/g of fresh weight. Low expression levels (2–4 mg/g of fresh weight) were also observed for His-tagged RBD variant (aa R319–F541, where the number of cysteine residues was not even) as reported by Diego-Martin et al. (2020) and Shin et al. (2021), which was too low to be economical for commercialization (*20,21*).

The plant produced VLP (virus–like particles, two doses of 3.75 μg of antigen) based vaccine has proven to show efficacy of 75.3% against COVID-19 (original Wuhan strain), however, there are no report about efficacy of this vaccine against Omicron. Because this plant-based VLP vaccine is targeted on the full-length S sequence, Medicago is preparing to study an Omicron-adapted version of its vaccine, as reported by D’Aoust (https://www.reuters.com/business/healthcare-pharmaceuticals/canada-approves-medicagos-plant-based-covid-19-vaccine-adults-2022-02-24/) even though Canada has already approved a plant-based VLP Medicago vaccine (designed on S protein of SARC-CoV-2 original Wuhan strain) for COVID-19 for adults.

In conclusion, based on the data we obtained, plant produced RBD and cocktail antigens are promising vaccine candidates against SARS-CoV-2, independently of its variants and may protect against Delta and Omicron-mediated COVID-19.

## Author Contributions

T.M. conceptualized the study; T.M. designed the experiments; D.Y., M.I., I.G., B.G., G.M., A.O., H.Y., B.K., S.T.I.P., M.A.U. performed the experiments; T.M. and G.H. analyzed the data; T.M. and G.H. contributed to writing the paper. All authors have read and agreed to the published version of the manuscript.

## Data and materials availability

All data are available in the main manuscript or the supplementary materials. The authors are grateful to George P. Lomonossoff (John Innes Center, Biological Chemistry Department) and Plant Bioscience Limited for kindly providing the pEAQ binary expression vector. We thank Philip de Leon at Trade Connections International for his editorial assistance.

